# Proteomic identification of phosphorylation dependent Septin 7 interactors that drive dendritic spine formation

**DOI:** 10.1101/2022.01.04.474970

**Authors:** Sujin Byeon, Bailey Werner, Reilly Falter, Kristian Davidsen, Calvin Snyder, Shao-En Ong, Smita Yadav

## Abstract

Septins are a family of cytoskeletal proteins that regulate several important aspects of neuronal development. Septin 7 (Sept7) is enriched at the base of dendritic spines in excitatory neurons and mediates both spine formation and spine-synapse maturation. Phosphorylation at a conserved C-terminal tail residue of Sept7 mediates its translocation into the dendritic spine head to allow spine-synapse maturation. The mechanistic basis for postsynaptic stability and compartmentalization conferred by phosphorylated Sept7, however, is not known. We report herein the proteomic identification of Sept7 phosphorylation dependent neuronal interactors. Using Sept7 C-terminal phosphopeptide pulldown and biochemical assays, we show that the 14-3-3 family of proteins specifically interact with Sept7 when phosphorylated at the T426 residue. Biochemically, we validate the interaction between Sept7 and 14-3-3 isoform gamma, and show that 14-3-3 gamma is also enriched in mature dendritic spine head. Further, we demonstrate that interaction of phosphorylated Sept7 with 14-3-3 protects it from dephosphorylation, as expression of a 14-3-3 antagonist significantly decreases phosphorylated Sept7 in neurons. This study identifies 14-3-3 proteins as an important physiological regulator of Sept7 function in neuronal development.

## INTRODUCTION

Septins are evolutionarily conserved cytoskeletal proteins important for diverse cellular processes including cell division, regulation of actin and microtubule dynamics, localization of scaffolding proteins, and membrane trafficking (Mostowy and Cossart, 2012). Septins structurally contain a GTP-binding domain and a variable N- and C-terminal domain which oligomerize with each other to form symmetric filaments, higher order structures such as filaments and rings (Mendonça et al., 2021; Sirajuddin et al., 2007). In humans, there are thirteen septin genes within four homology groups: *SEPT2* (*SEPT1, SEPT2, SEPT4*, and *SEPT5*), *SEPT3* (*SEPT3, SEPT9*, and *SEPT12*), *SEPT6* (*SEPT6, SEPT8, SEPT10, SEPT11*, and *SEPT14*), and *SEPT7* (*SEPT7*) (Kinoshita, 2003a). Within these septin oligomers, septins from the same group are interchangeable, suggesting potential redundancy in their function. Notably, forming its own group, Septin7 (Sept7) is a unique and non-redundant core component of the septin complexes (Kinoshita, 2003b).

Several members of the septin family were found to be enriched in the postsynaptic density (PSD) fractions from mouse brain by mass-spectrometry analysis, with Sept7 suggested to be the most abundant (Walikonis et al., 2000). Sept7 is expressed throughout all stages of neuronal differentiation and localizes at axonal and dendritic branching points as well as at the base of the dendritic protrusions (Tada et al., 2007; Xie et al., 2007). Depletion of Sept7 leads to decreased branching of axon and dendrites both *in vitro* and *in vivo* (Ageta-Ishihara et al., 2013; Hu et al., 2012; Tada et al., 2007; Xie et al., 2007). Moreover, an increase in the number of immature dendritic filopodia were observed when Sept7 was knocked down (Tada et al., 2007). Sept7 localizes at the base of dendritic spines to create an important diffusion barrier for membrane protein entry into the dendritic spine head (Ewers et al., 2014). Further, Sept7 was found to be a phosphorylation target of the autism risk gene *TAOK2* which encodes a serine threonine kinase (Anda et al., 2012; Nourbakhsh et al., 2021; Richter et al., 2018; Yadav et al., 2017). Phosphorylation of Sept7 by TAOK2 at an evolutionarily conserved C-terminal tail residue T426 was shown to be essential for spine maturation. While in its unphosphorylated state Sept7 localizes to the base of dendritic spines and filopodia, Sept7 when phosphorylated at T426, relocates to the dendritic spine head (Yadav et al., 2017). Although mechanisms through which phosphorylation induces Sept7 translocation were unclear, it was found that preventing this phosphorylation leads to increased dendritic filopodia as well as mislocalization of the synaptic scaffold proteins to the dendritic shaft instead of the spine. This eventually led to mislocalized synapses that formed on the dendritic shafts (Yadav et al., 2017). How phosphorylation at the conserved T426 residue on the C-terminal tail of Sept7 regulates its function is unknown. The C-terminal tail extends perpendicularly out to the axis of the oligomeric septin filament (Sirajuddin et al., 2007). This is thought to facilitate protein-protein interactions in addition to lateral associations between septin filaments (Finnigan et al., 2015, 2016; Marques et al., 2012).

Herein, we show that phosphorylation of Sept7 extended C-terminal tail at T426 residue can be mediated by several members of the TAO kinase family including TAOK1 and two distinct isoforms of TAOK2 kinase. Further, Sept7 phosphorylation at T426 was found to increase during development in rat embryonic hippocampal neurons. Expression of phosphomimetic Sept7 T426D leads to earlier maturation of dendritic spines in these neurons. Using live confocal imaging, we show that phosphorylated Sept7 is stably associated with the dendritic spine-head over time in contrast to the phosphomutant Sept7. To identify potential mechanisms through which phosphorylated Sept7 contributes to dendritic spine maturation, we performed an unbiased proteomic screen to identify phosphorylation dependent binding partners of Sept7. Among the identified candidate interactors were several isoforms of 14-3-3 proteins and actin binding proteins. Since 14-3-3 proteins bind and modulate functions of phosphorylated proteins, we tested whether phosphorylated Sept7 associates with 14-3-3 proteins. Using biochemical assays, we demonstrate that phosphorylated Sept7 interacts with 14-3-3 gamma isoform. Further, we found that 14-3-3γ gamma is enriched in dendritic spines compared to other 14-3-3 isoforms. Finally, we found that disrupting 14-3-3 and Sept7 interaction perturbs level of phosphorylated Sept7 in neurons, revealing an important role of this interaction in regulating neuronal development.

## MATERIALS AND METHODS

### Antibodies and Plasmids

Antibodies used for these experiments include: anti-HA-tag (Mouse, ProteinTech, Thermo Fisher, 50-173-6449), phospho-TAOK2 (S181) (Rabbit, R&D Systems, PPS037), GFP (Mouse, Roche, 11814460001), GST (Mouse, Invitrogen, MA4-004). Rabbit phospho-Septin7 (pT426) antibody was generated as described previously (Yadav et al., 2017). Human TAOK2α was PCR amplified from pCMV-Sp6-TAOK2 plasmid (Ultanir et al., 2014) and cloned into the sfGFP-C1 vector (Addgene #54579) using restriction sites HindIII and MfeI. Human TAOK2β cDNA was obtained from Transomics (#BC152413) and cloned into the sfGFP-C1 vector using the same restriction sites. Similarly, TAOK1 cDNA from Transomics (#BC144067) was inserted into the sfGFP-C1 vector using restriction enzymes. Addgene plasmids used for these experiments include: HA-14-3-3 zeta (#116888), pcDNA3-HA-14-3-3 gamma (#13274). EYFP-Difopein and EYFP-Control were gifts from Dr. Yi Zhou (Florida State University).

### Immunoprecipitation Kinase Assays

sfGFP tagged TAOK1, TAOK2α, and TAOK2β constructs (3μg) were expressed in HEK293T cells grown in Dulbecco’s Modified Eagle Medium (DMEM) containing 10% Fetal Bovine Serum (FBS) and 1% Penicillin-Streptomycin. 24 hours post transfection, cells were collected and incubated in HKT lysis buffer (25mM HEPES pH 7.2, 150mM KCl, 1% Triton X-100, 2mM DTT, 1x Protease Inhibitor (Roche, cOmplete™ EDTA free)) for 30 minutes on ice prior to homogenization with a 25-gauge syringe needle. Pierce™ Protein G Agarose beads (Thermo Fisher) were washed with HKT buffer three times. The supernatant was collected from the cell lysates through centrifugation at 800g for 5 minutes, precleared with Protein G Agarose beads for 30 minutes, and immunoprecipitated with the GFP antibody overnight at 4°C. Beads were washed with HKT buffer twice, incubated with high salt HKT buffer (25mM HEPES pH 7.2, 1M NaCl, 1% Triton X-100, 2mM DTT, 1x Protease Inhibitor (Roche, cOmplete™ EDTA free)) for 10 minutes, washed with HK buffer (25mM HEPES, 150mM KCl, 2mM DTT, 1x Protease Inhibitor (Roche, cOmplete™ EDTA free)) twice and then with the kinase buffer (20mM Tris HCl pH 7.5, 10mM MgCl2, 1mM DTT, 1x Protease Inhibitor (Roche, cOmplete™ EDTA free)) twice at 4°C. Kinase assay was performed by incubating the beads with the kinase buffer, 1mM ATP, and 10x Phosphatase Inhibitor (ThermoFisher, Halt) at 30°C for 45 minutes at 920rpm. To assay for phosphorylation of Septin7 by TAO kinases, purified GST tagged Septin7 C-terminal tail (321-438 amino acids) was also added to the reaction. Samples were prepared by adding NuPAGE™ LDS Sample Buffer (Thermo Fisher) containing 125mM DTT, heating for 10 minutes at 95°C, and centrifuging at 5,000g for 5 minutes. Samples were run on NuPAGE™ 4-12% Bis-Tris Polyacrylamide gels (Thermo Fisher) with NuPAGE™ MOPS running buffer (Thermo Fisher) at 165V for 20 minutes and then 175V for 50 minutes. Gels were transferred to Immobilon-P membrane at 100V for 1 hour. The blots were blocked with 5% BSA blocking buffer and probed with Phospho-TAO2 (S181) and Phospho-Septin7 (T426) antibodies at 1:500 dilution overnight at 4°C followed by 3 hour incubation with HRP conjugated secondary antibody at 1:5000 dilution at room temperature. The kinase activity was quantified by normalizing Phospho-TAO2 signal intensity to that of GFP signal and Phospho-Septin7 signal intensity to that of GST signal.

### Co-immunoprecipitation

Co-immunoprecipitation assays were performed in HEK293T cells grown in DMEM media (Thermo Fisher, Gibco) with 10% fetal bovine serum (Axenia) and 1% Penicillin-Streptomycin (Invitrogen). Cells were grown to confluence in a flat-bottom plate with 35 mm wells at 5% CO2 and 37°C and transfected with 2.0μg of HA-14-3-3 gamma, HA-14-3-3 zeta, or pRK5-HA. Approximately 36 hours post-transfection, cells were treated with 0.1μM Okadaic acid for 10 minutes and lysed with 1 mL HKT buffer (25mM HEPES pH7.2, 100mM KCl, 1% Triton X-100, 1mM DTT, 1 mM EDTA, Phosphatase Inhibitor (Thermo Fisher, Halt) Protease Inhibitor (Roche, cOmplete™ EDTA free)) per 2-well construct. Lysate was incubated on ice for 20 minutes prior to homogenization with a 25-gauge syringe needle. Homogenate was pelleted via centrifugation at 6,000g and 4°C for 5 minutes. Pierce™ ProteinG Agarose (Thermo Fisher) beads and EZview™ Red Anti-HA Affinity Gel (Millipore-Sigma) beads were washed twice for 5 minutes at 4°C and re-suspended in 150μL HKT per sample tube. Pellet supernatant was pre-cleared with 20μl Pierce™ ProteinG Agarose (Thermo Fisher) beads, and pre-cleared supernatant was collected for input samples. Remaining volume of pre-cleared supernatant was immunoprecipitated with 20μl EZview™ Red Anti-HA Affinity Gel (Millipore-Sigma) beads. Beads were washed three times with HKT and twice with HK buffer (25mM HEPES pH 7.2, 100mM KCl, 1mM DTT, 1 mM EDTA). Both immunoprecipitation and input samples were prepared with 4X LDS Sample Buffer (Thermo Fisher) and 125 mM DTT followed by heat treatment at 95°C for 10 minutes. Samples were centrifuged at 14,000g for 5 minutes at 4°C and electrophoresed on NuPAGE™ 4-12% Bis-Tris Polyacrylamide gels (Thermo Fisher) with NuPAGE™ MOPS running buffer (Thermo Fisher) for 40 minutes at 160V.

### Western Blot

For western blot analysis, gels were transferred to ImmobilonP PVDF membrane (Millipore-Sigma) with transfer buffer (25mM Tris, 192mM Glycine, 20% (v/v) Methanol) for 60 minutes at 100 V. Blots were cut below the 50 kDa mark to produce one half-membrane consisting of high molecular weights to detect phosphorylated Septin-7 and one half-membrane consisting of low molecular weights to detect HA-tagged 14-3-3 isoforms. Blots were blocked in 5% milk (5% milk powder (Carnation), 50mM Tris-Cl, 150mM NaCl, 0.1% Tween™20 (Thermo Fisher)) or 2% BSA (2% Bovine Serum Albumin (VWR), 50mM Tris-Cl, 150mM NaCl, 0.1% Tween™20 (Thermo Fisher)) blocking buffer for one hour at room temperature and incubated in primary antibody overnight at 4°C. Primary rabbit antibody to detect phosphorylated Septin-7 (T426) was diluted to 1:250 and applied to high molecular weight half-membrane, and primary mouse antibody to detect HA-tag (ProteinTech) was diluted to 1:1000 and applied to low molecular weight half-membrane. Blots were washed with blocking buffer at room temperature before secondary antibody incubation. Blots were incubated in HRP-conjugated secondary antibody at 1:1000 dilution for three hours at room temperature. Blots were washed successively with blocking buffer, TBST (50mM Tris-Cl, 150mM NaCl, 0.1% Tween™20 (Thermo Fisher)), and TBS (50mM Tris-Cl, 150mM NaCl). Western blot images were visualization with Pierce™ ECL Western Blotting Substrate (Thermo Fisher) and captured with the ChemiDoc Imager (BioRad).

### Peptide Pulldown and Mass Spectrometry

Synthetic peptide corresponding to Sept7 C-terminal tail residues 416-438 were commercially synthesized with an N-terminal Cysteine residue and T426 in either phosphorylated or unphosphorylated forms (Elim Biopharmaceuticals). Peptides were coupled to SulfoLink Beads (Invitrogen) according to manufacturer’s protocol. Briefly, 0.5mg of peptide was dissolved in 1ml of coupling buffer (50mM Tris and 5mM EDTA, pH8.5) and then TCEP was added to a final concentration of 25mM. Beads (50ul for each peptide) were washed 4 times in the coupling buffer, quenched with L-cysteine-HCl and then incubated with 100ul of 0.5mg/ml P15 mouse brain lysate. Beads were washed 4 times with lysis buffer containing 20mM Tris pH 7.5, 100mM NaCl, 10mM MgCl2, 0.5mM DTT, 1% Triton X-100, 0.1% deoxycholic acid and protease inhibitors (Roche), and thrice with lysis buffer without detergent. Proteins bound to beads were then denatured by adding 3x bead volumes of 8M Urea/10 mM Tris pH 8.0 to resuspend beads. TCEP was added to a final concentration of 1mM and incubated at room temperature for 30 min with thermomixer/slow vortex. Chloroacetamide (CAM) was added to final concentration of 3mM and incubated at room temperature for 10 min with thermomixer/slow vortex. TCEP was added to quench excess CAM after alkylation is complete. The pH was then adjusted to 8. For protein digestion LysC (Mass Spec Grade) at 1:100 enzyme:substrate ratio and added and incubated at 37°C for 2hr on a thermomixer with gentle agitation. Then 3x reaction volumes of TEAB was added to dilute urea to less than or equal 2M Urea. Then pH was adjusted to 8.0. Trypsin (MS grade, Promega) was added at 1:100 enzyme:substrate ratio and incubated at 37°C for 12-16hr with gentle agitation. Digestion is stopped by adding TFA to a final concentration of 1%. Peptide samples were desalted on C18 StageTips. Peptide samples were separated on an Thermo Dionex Ultimate 3000 RSLCnano System (Thermo Fisher Scientific) using 20 cm long fused silica capillary columns (100 *μ*m ID, laser pulled in-house with Sutter P-2000, Novato CA) packed with 3 μm 120 Å reversed phase C18 beads (Dr. Maisch, Ammerbuch, DE). Liquid chromatography (LC) solvent A was 0.1% (v/v) aq. acetic acid and LC solvent B was 20% 0.1% (v/v) acetic acid, 80% acetonitrile. The LC gradient was 100 min long with 5-35% B at 300 nL/min. MS data was collected with a Thermo Fisher Scientific Orbitrap Elite. Data-dependent analysis was applied using Top15 selection with CID fragmentation. Data .raw files were analyzed by MaxQuant/Andromeda (Cox et al., 2011) version 1.6.2.8 using protein, peptide and site FDRs of 0.01 and a score minimum of 40 for modified peptides, 0 for unmodified peptides; delta score minimum of 17 for modified peptides, 0 for unmodified peptides. MS/MS spectra were searched against the UniProt mouse database (updated July 2016). MaxQuant search parameters: Variable modifications included Oxidation (M) and Phospho (S/T/Y). Carbamidomethyl (C) was a fixed modification. Max. missed cleavages was 2, enzyme was Trypsin/P and max. charge was 7. The initial search tolerance for FTMS scans was 20 ppm and 0.5 Da for ITMS MS/MS scans.

### Immunofluorescence

Rat embryonic hippocampal neurons were grown on coverslips coated with 0.1M borate buffer (50mM boric acid, 12.5mM sodium tetraborate in water, pH 8.5) 60ug/mL Poly-D-Lysine (Sigma), and 2.5ug/mL Laminin (Sigma) in 12 well plates. For immunofluorescence, neurons were fixed with warm 4% paraformaldehyde + 4% sucrose in PBS, followed by 60 min incubation in blocking buffer (0.2% Triton X-100, 10% Normal Goat Serum (Jackson Labs), and 0.2M Glycine pH 7.2) at room temperature. Primary antibody incubation was overnight at 4°C followed by secondary antibody for three hours at room temperature. Coverslips were mounted on slides using Fluoromont-G.

### Neuronal Culture

Hippocampi were obtained from E17-E18 Sprague-Dawley rat embryos (Envigo), trypsin dissociated and plated at a density of 150,000 cells per 18mm coverslip (Fisher) and 50,000-500,000 cells per 35mm glass bottom dish (MatTek). Dishes were coated as previously described. Neurons were seeded in plating media (10% Fetal bovine serum heat inactivated, 20% dextrose, 1x Glutamax (Invitrogen), 1x penicillin/streptomycin, MEM Eagle’s with Earle’s BBS (Lonza)) for four hours. Media was changed to maintenance media (B27 (Invitrogen), 1x penicillin/streptomycin, 1x Glutamax, Neurobasal media (Invitrogen)). Half the media was replaced with new maintenance media every 3-4 days.

### Microscopy and Image Analyses

All live and fixed cell imaging was performed on a Nikon Ti2 Eclipse-CSU-X1 confocal spinning disk microscope equipped with four laser lines 405nm, 488nm, 561nm and 670nm and an sCMOS Andor camera for image acquisition. The microscope was caged within the OkoLab environmental control setup enabling temperature and CO2 control during live imaging. Imaging was performed using Nikon 1.49 100x Apo 60X or 40X oil objectives. All image analyses were done using the open access Fiji software.

### Statistics

All statistics except for the mass spectrometry data were performed in GraphPad software Prism9.0. Two group comparisons were made using unpaired t-test unless otherwise stated. Statistically, *p* value less that 0.05 was considered significant. All experiments were done in triplicate unless stated otherwise, and experimental sample size and *p* values are indicated with the corresponding figures.

## RESULTS

### The conserved C-terminal tail in Septin 7 is phosphorylated by TAO kinases during neuronal development

Sept7 was identified through an unbiased chemical-genetic screen as a phosphorylation target of TAOK2 (Yadav et al., 2017). The site of phosphorylation residue T426 lies in the extended C-terminal coiled coil tail of Sept7. This threonine residue which harbors the consensus site for TAO kinases p[S/T]-X-X[R/H/K] is evolutionarily highly conserved (Figure 1A). Given the structural conservation among the kinase domain of TAO kinases, we tested whether closely paralogous kinase TAOK1 and the alternatively spliced isoforms of TAOK2, TAOK2α and TAOK2β could phosphorylate Sept7 at T426. GFP tagged TAOK1, TAOK2α and TAOK2β were expressed independently in HEK293T cells, immunoprecipitated using GFP antibody and incubated with purified GST-Septin7 protein (amino acids 321-428) in a kinase reaction. Using a phosphospecific antibody (pT426) against Sept7 T426 residue, we found that TAOK1, TAOK2α and TAOK2β could each phosphorylate purified GST-Sept7 (Figure 1B and 1C). Further, by immunostaining cultured hippocampal neurons at different stages of development, we found that phosphorylated Sept7 (T426) levels increased in neurons as it matured from DIV4, DIV9 to DIV14 (Figure 1D, E). This is consistent with the increased levels of TAOK2 expression in developing neurons (Anda et al., 2012).

**Figure 1.**
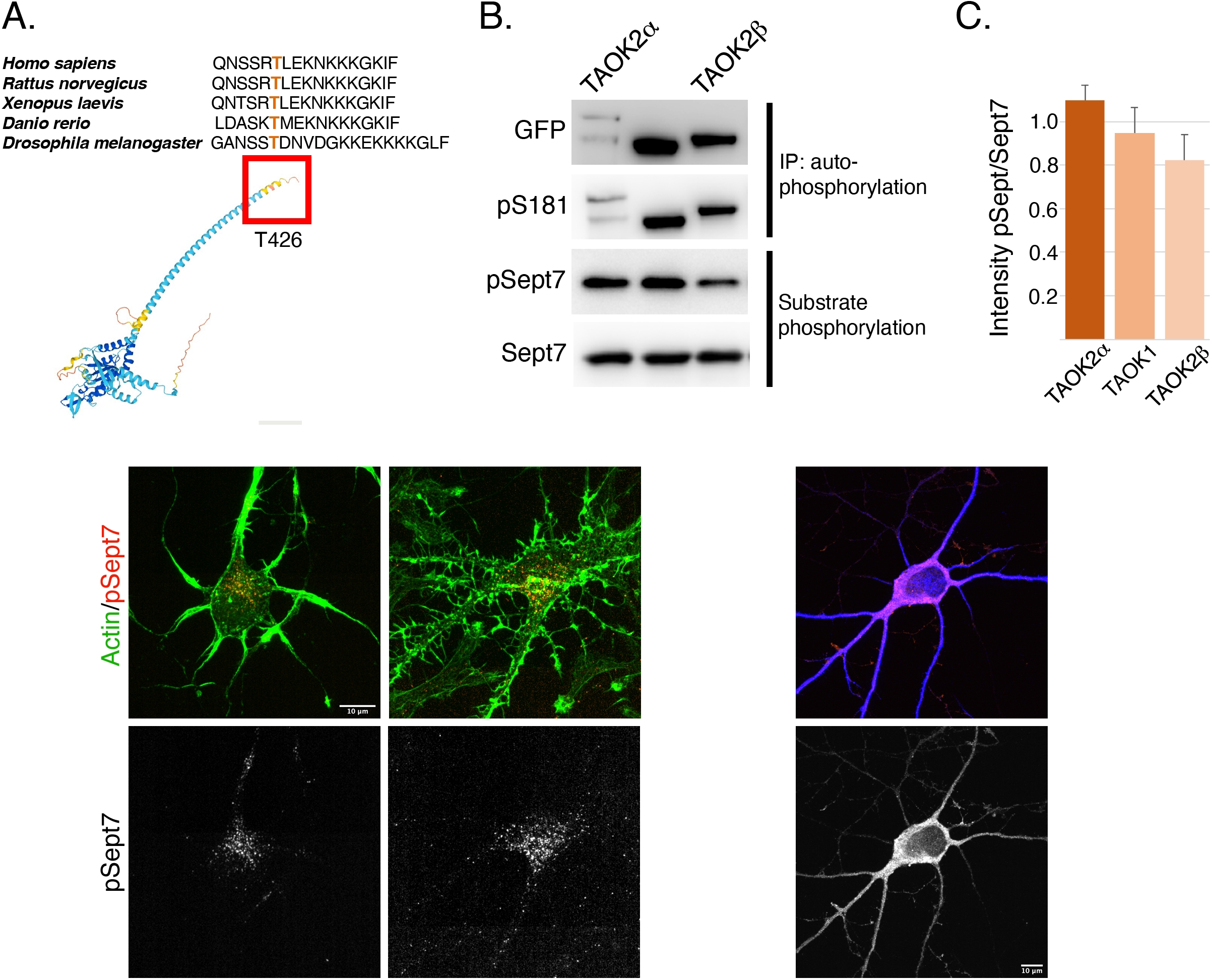
TAO kinases phosphorylate the C-terminal tail of Sept7 during neuronal development. (A) Top: Multiple sequence alignment of the C-terminal tail of Sept7 demonstrating it evolutionarily conserved sequence including the phosphorylation site. The T426 site of phosphorylation is shown in orange. Bottom: AlphaFold2.0 predicted protein structure of Sept7 in which the T426 phosphorylation site is outlined in red. (B) Western blot shows kinase activity of sfGFP-TAOK2α, sfGFP-TAOK1, or sfGFP-TAOK2β expressed in HEK293T cells, immunoprecipitated with GFP antibody, and incubated with purified GST-Sept7 C-terminal tail (321-438 amino acids) for kinase reaction. Substrate phosphorylation was determined by probing for phosphorylated Sept7 using the pT426 antibody. Total Sept7 was detected by antibody against GST tag. TAO kinase autophosphorylation was determined by probing for phosphorylated S181 residue. (C) Quantification of protein band intensity of western blot in (B). Bar graph displays the ratio of phosphorylated Sept7 band intensity to GST band intensity for each TAO kinase construct, in which no significant difference was observed between TAO kinases. Error bars represent standard deviation from n=2 experiments. (D) Fixed images of DIV4 and DIV9 hippocampal neurons immunostained for endogenous phosphorylated Sept7 T426 and phalloidin to visualize actin. Scale bar represents 10μm. (E) Fixed images of DIV14 hippocampal neurons immunostained for endogenous phosphorylated Sept7 T426 and dendritic marker MAP2. Scale bar represents 10μm.

### Altered dynamics of phosphorylated Septin 7

We had previously reported that phosphorylated Sept7 is enriched in the dendritic spine-head and is important for dendritic spine formation and stability of PSD95 (Yadav et al., 2017). Expression of Sept7 phosphomutant T426A leads to failure of dendritic spine maturation, resulting in exuberant dendritic filopodia and shaft synapses (Yadav et al., 2017). To test the role of Sept7 phosphorylation during neuronal maturation, we expressed phosphomimetic (T426D) and phosphomutant (T426A) Sept7 in DIV9 neurons, a developmental stage where filopodial protrusions have not yet developed into mature mushroom dendritic spines. We found that expression of GFP-tagged phosphomimetic Sept7 (T426D) as opposed to phosphomutant Sept7 (T426A) led to early maturation of dendritic spines in DIV9 hippocampal neurons (Figure 2A). Further, there was accumulation of Sept7 in the dendritic spines in neurons expressing phosphomimetic Sept7 T426D, where 86.2% of spines were positive for the expressed GFP-Sept7 (Figure 2C). To determine how phosphorylation at its C-terminal tail residue T426 affects Sept7 dynamics, we performed confocal live imaging of neurons expressing either the phosphomimetic (T426D) or phosphomutant (T426A) GFP-tagged Sept7. Using live imaging, we found that phosphomimetic Sept7 (T426D) remained stably localized within the dendritic spine head, while the phosphomutant Sept7 (T426A) predominantly was localized at the base of the dendritic spine (Figure 2B, montage over 30sec). Based on these data showing distinct localization and dynamics of Sept7 in its phosphorylated and unphosphorylated states at residue T426, we hypothesized that there must be unique interaction partners of phosphorylated Sept7 that modulate its discrete properties.

**Figure 2.**
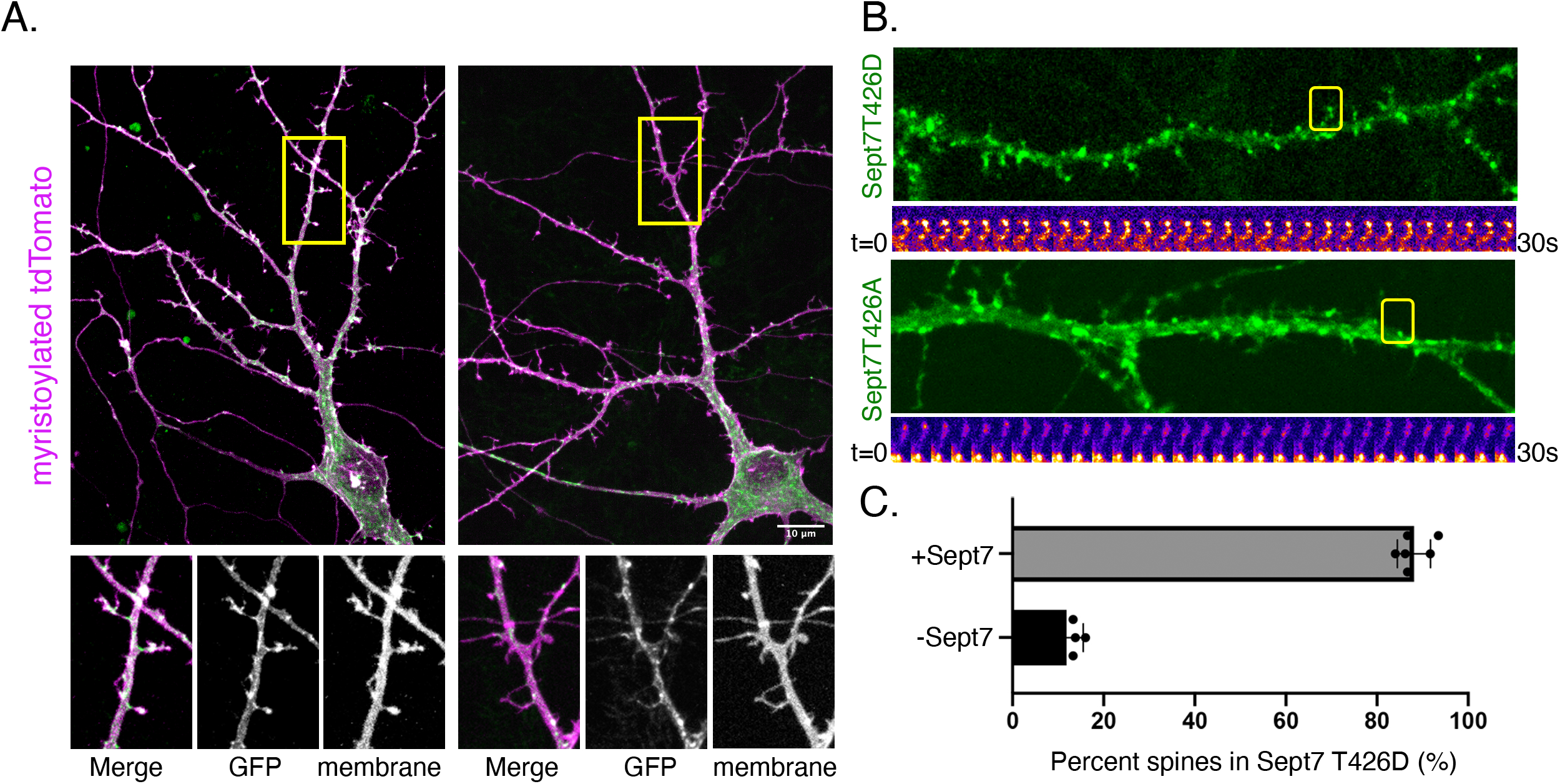
Phosphorylation of Sept7 alters its localization and dynamics. (A) Fixed images of DIV9 hippocampal neurons transfected with phosphomimetic GFP-Sept7(T426D) or phosphomutant GFP-Sept7(T426A) along with membrane marker myristoylated-tdTomato. Phosphomimetic GFP-Sept7 (T426D) expression leads to early maturation of dendritic spines compared to phosphomutant GFP-Sept7 (T426A). (B) Live imaging montage over 30sec of DIV9 hippocampal neurons transfected with phosphomimetic GFP-Sept7(T426D) or phosphomutant GFP-Sept7(T426A). Phosphomimetic GFP-Sept7 (T426D) localizes to the dendritic spine head, whereas phosphomutant GFP-Sept7 (T426A) localizes to the base of the dendritic spine. (C) Quantification of percent of dendritic spines positive or negative for Sept7 in DIV9 hippocampal neurons transfected with phosphomimetic GFP-Sept7(T426D). Sept7 accumulates in dendritic spines in neurons expressing phosphomimetic GFP-Sept7 (T426D). Values represent mean, error bars represent SEM and n=6.

### Proteomic identification of Septin 7 phosphorylation dependent binding partners

We set out to identify proteins that specifically interact with Sept7 following phosphorylation at residue T426 in its C-terminal tail. This was achieved using an unbiased proteomic strategy to pull down proteins from P15 mouse brain lysate with a synthetic peptide corresponding to Sept7 C-terminal tail residues 416-438 (Figure 3A). This peptide of 22 amino acids was unique to Sept7, as BLAST analyses did not identify any other protein harboring this sequence. Next, the phosphorylated (pSept7) and non-phosphorylated (Sept7) peptides were coupled to beads. Agarose beads activated with iodoacetamide were used for covalent immobilization of peptides corresponding to pSept7 and Sept7 peptides. Peptide coupled beads were then incubated with mouse brain lysate. Proteins that were bound to peptide-beads were enzymatically digested, and then subjected to mass spectrometry-based identification. Label free quantification was used to measure the relative abundance of proteins bound to Sept7 and pT426-Sept7 (Figure 3A). In our analyses of 5 replicates of peptide pulldowns, we found 37 proteins that were significantly associated with phosphorylated Septin7 peptide over unphosphorylated peptide (Figure 3B). These proteins were analyzed by String11.2 and the most significant biological process was identified as regulation of cytoskeletal organization (Figure 3C, E) including actin binding proteins as well the phosphoprotein regulating proteins of the 14-3-3 family (Figure 3D, E). The 14-3-3 protein family is a group of highly conserved acidic proteins highly expressed in the mammalian brain (Ferl et al. 2002, Boston et al. 1982). In humans, there are seven known isoforms of 14-3-3: β, γ, ε, η, δ, τ, and ζ (Gardino et al. 2006). Forming both homo- and hetero-dimers, 14-3-3 proteins are ubiquitous regulatory proteins that can simultaneously bind one or two client proteins with their individual ligand-binding channels (Gardino et al. 2006, Uhart & Bustos 2013). Among the candidate pSept7 binding partners identified by our mass spectrometry experiment (Figure 3C), we were specifically interested in 14-3-3 proteins as 1) binding of 14-3-3 proteins is largely determined by phosphorylation of the client, with the isoforms typically binding phosphorylated serine/threonine motifs (Fu et al. 2000, Aitken 1996), 2) 14-3-3 are expressed in the brain and important for synapse development, and 3) binding of phosphoproteins to 14-3-3 can serve diverse purposes, including preventing protein-protein interactions, scaffolding, inducing conformational changes, protecting phosphorylation sites, and promoting or preventing ubiquitination.

**Figure 3.**
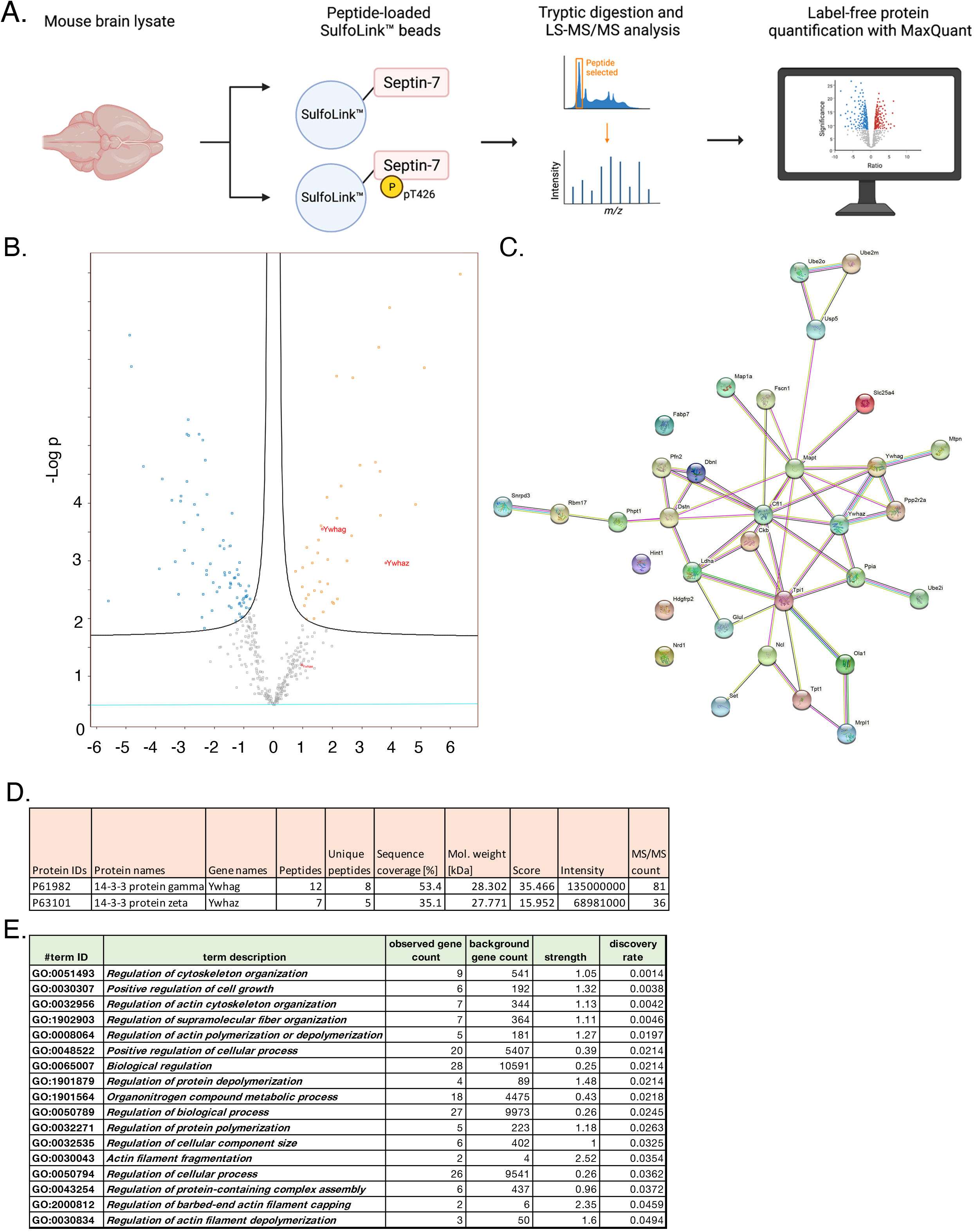
Proteomic identification of Septin 7 phosphorylation-dependent binding partners. (A) Schematic representation of the proteomic strategy for identifying Sept7 phosphorylation-dependent binding partners. (B) Proteomic results of strategy described in (A). Volcano plot displays the difference in association of proteins with phosphorylated Sept7 versus unphosphorylated Sept7 based on 5 replicate pulldowns. 37 proteins were significantly associated with phosphorylated Sept7 peptide. 14-3-3 protein isoforms are labeled in red. (C) Protein-protein association network (STRING v11) of candidate pSept7 interaction partners. (D) Mass spectrometry results showing intensities for 14-3-3 protein isoforms gamma and zeta, which were specifically associated with phosphorylated Sept7 but not Sept7. (E) Biological processes associated with the protein-protein association network in (C) as identified through STRING v11.

### 14-3-3 proteins associate with Septin 7

We next validated whether 14-3-3 proteins were *bona fide* interacting proteins of Sept7 using co-immunoprecipitation assays. Using an antibody that recognizes all isoforms of 14-3-3 (pan 14-3-3 polyclonal), we immunoprecipitated 14-3-3 proteins from neuronal lysate obtained from DIV18 cultured embryonic rat hippocampal neurons, and then probed for phosphorylated Sept7 using the pT426 Sept7 antibody. We found that indeed pSept7 coimmunoprecipitated with 14-3-3 proteins from neuronal lysates (Figure 4A). Since 14-3-3 zeta (ζ) and gamma (γ) were the isoforms most enriched in our proteomic data (Figure 3D), and are also highly expressed in the brain (Berg et al., 2003), we next tested if there was an interaction of these isoforms with phosphorylated Sept7. We co-expressed HEK293T cells with GFP-Sept7 along with either HA tagged 14-3-3 zeta, 14-3-3 γ, or both isoforms together. Using the HA antibody, we pulled down the 14-3-3 isoforms and then probed for phosphorylated Sept7 using the pT426 antibody. We found that pSept7 co-immunoprecipitated strongly with 14-3-3 γ, and 14-3-3(ζ + γ) but not 14-3-3ζ (Figure 4B). Further, in order to determine the effect of expression of different 14-3-3 isoforms on dendritic spine maturation, we co-expressed HA-tagged 14-3-3γ and 14-3-3ζ along with membrane marker myr-tdTomato in cultured DIV12 hippocampal neurons. Neurons were fixed at DIV14 and stained with anti-HA antibody to visualize the localization of 14-3-3 proteins (Figure 4C). In neurons expressing HA-14-3-3γ, about 52% of protrusions were spiny mushroom mature spines, while 42% of protrusions were spiny in neurons expressing HA-14-3-3ζ (Figure 4D). Interestingly, HA-14-3-3γ was enriched in dendritic spines with a spine/dendrite ratio of 1.2, whereas HA-14-3-3ζ primarily localized to the dendritic shaft and was not enriched in the spine head (Figure 4E). In summary, we found that phosphorylated Sept7 interacts with 14-3-3γ, and that both pSept7 and 14-3-3γ are enriched in the dendritic spine head and enhance spine maturation.

**Figure 4.**
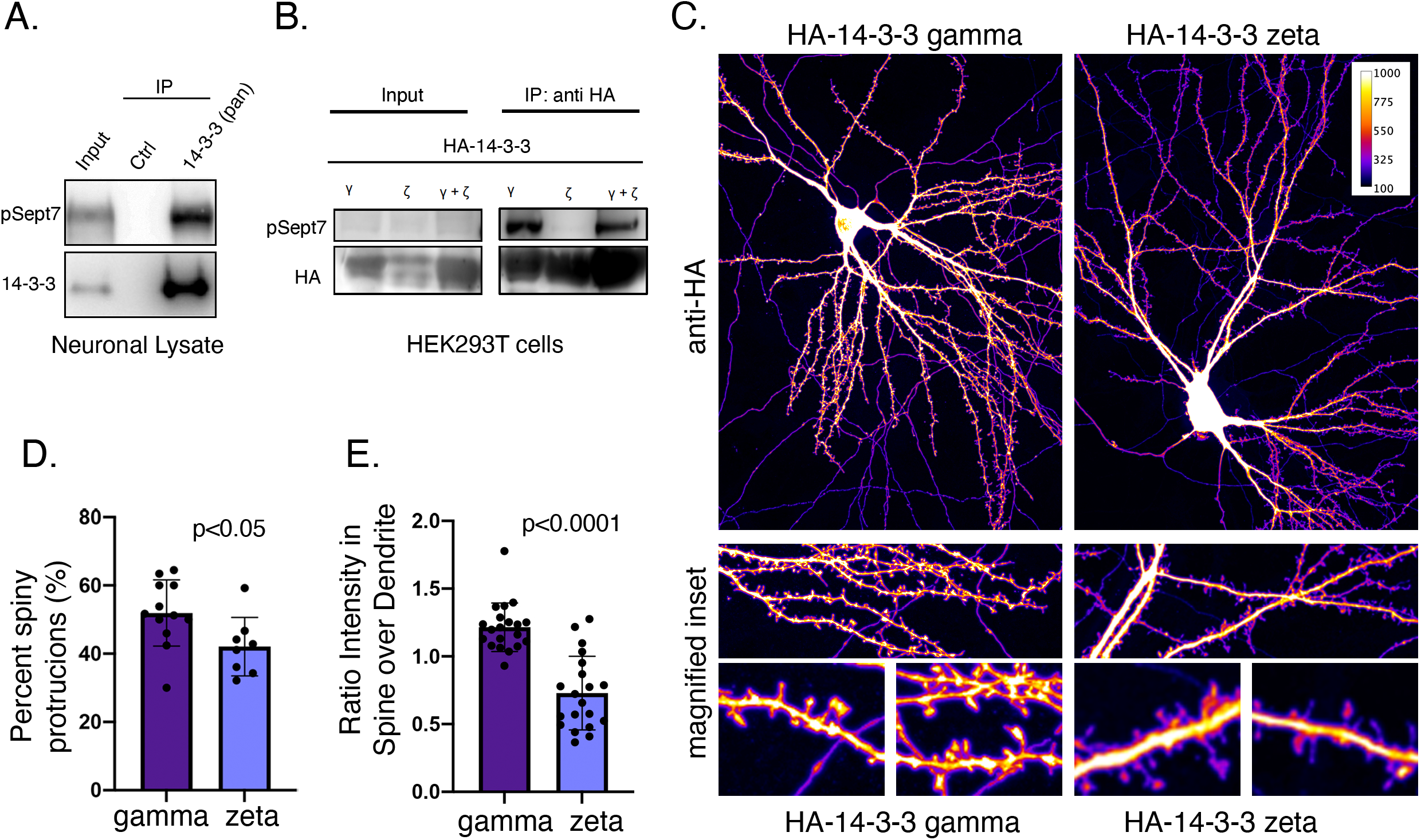
14-3-3γ associates with phosphorylated Sept7 and enhances dendritic spine maturation. (A) Western blot of endogenous 14-3-3 protein in DIV18 hippocampal neurons immunoprecipitated with 14-3-3 (pan) antibody and probed for 14-3-3 and phosphorylated Sept7. (B) Western blot of HA-14-3-3γ, HA-14-3-3ζ, or both HA-tagged isoforms co-expressed along with GFP-Sept7 in HEK293T cells immunoprecipitated with HA antibody and probed for HA and phosphorylated Sept7. Phosphorylated Sept7 co-immunoprecipitated with HA-14-3-3γ and HA-14-3-3(ζ+γ) but not HA-14-3-3ζ. (C) Fixed images of DIV14 hippocampal neurons co-transfected with membrane marker myristoylated tdTomato and HA-14-3-3γ or HA-14-3-3ζ and immunostained for HA tag. (D) Bar graph depicts percent of protrusions with mature, spiny morphology in DIV14 hippocampal neurons transfected with HA-14-3-3γ or HA-14-3-3ζ. HA-14-3-3γ expression results in significantly more spiny, mature protrusions (p<0.05). Values indicate mean, error bars represent SEM and n=5 neurons each. (E) Bar graph shows ratio of 14-3-3 fluorescent intensity in spines to intensity in corresponding dendritic shaft in DIV14 hippocampal neurons transfected with HA-14-3-3γ or HA-14-3-3ζ. Values indicate mean, error bars represent SEM, and n=20 spines from 5 neurons each.

### Sept7 association with 14-3-3 protects its phosphorylation at T426

To test the functional consequence of 14-3-3 association with phosphorylated Sept7, we utilized a genetically expressed inhibitor of 14-3-3, EYFP-Difopein (dimeric fourteen-three-three peptide inhibitor). This construct EYFP-Difopein is based on R18, a Raf-1-derived phosphopeptide, which is a high-affinity peptide antagonist of 14-3-3 proteins (Wang et al., 1999). Difopein is composed of two R18 coding sequences separated by a sequence coding for a short peptide linker in an EYFP fusion mammalian vector. Expressed Difopein is capable of disrupting 14-3-3/ligand binding (Masters and Fu, 2001). Difopein binds 14-3-3 without any isoform selectivity and thereby inhibits the interaction of all 14-3-3 isoforms with their physiological binding partners (Qiao et al., 2014). We transfected DIV7 hippocampal neurons with either control EYFP construct or EYFP-Difopein construct and then measured the levels of pSept7 in neurons at DIV9 (Figure 5A). Cells expressing Difopein had a significant decrease in the level of phosphorylated Sept7 compared to those expressing EYFP control, as detected by immunostaining using the pT426 Sept7 antibody (Figure 5A, yellow arrows and 5B). These data suggest that interaction of the C-terminal tail of Sept7 with 14-3-3 proteins is protective of the phosphorylation status of Sept7 in neurons, revealing a mechanism for modulation of Sept7 function by association with 14-3-3 proteins (Figure 5C).

**Figure 5.**
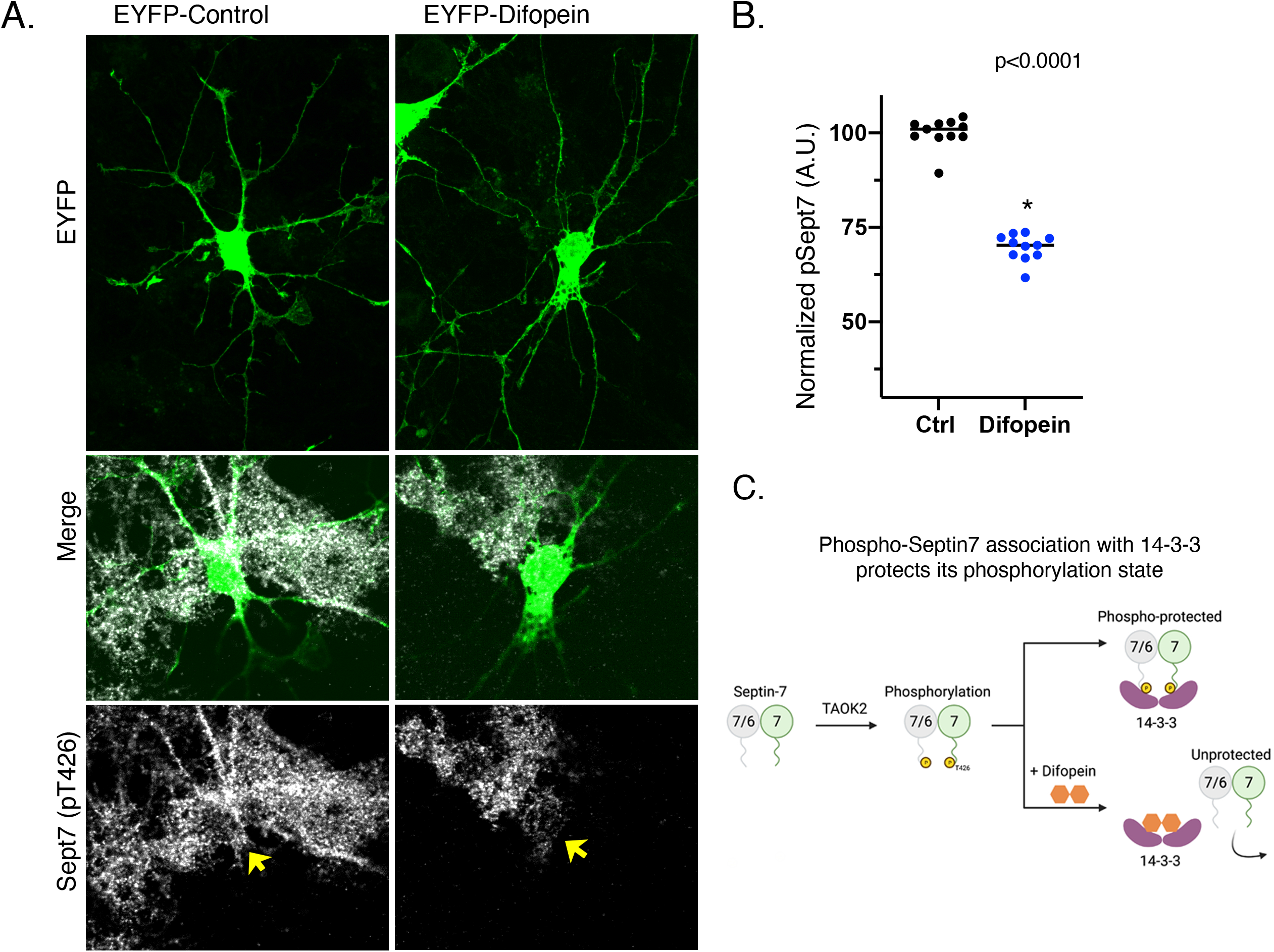
Association with 14-3-3 protects phosphorylation state of Sept7. (A) Fixed images of DIV9 hippocampal neurons transfected with EYFP-control or EYFP-Difopein, and immunostained for phosphorylated Sept7 (pT426). Phosphorylated Sept7 level was decreased in the difopein-expressing neuron (yellow arrows). (B) Quantification of phosphorylated Sept7 levels in the soma of DIV9 hippocampal neurons expressing EYFP-control or EYFP-Difopein. EYFP-Difopein expression results in significantly decreased levels of phosphorylated Sept7 (p<0.0001). Values indicate mean, error bars represent SEM and n=11 neurons each condition. (C) Schematic summarizes findings of this study showing 14-3-3 protein association modulates Sept7 function in neurons through protection of its phosphorylation state.

## DISCUSSION

In this study, we utilize mass spectrometry based discovery proteomics to identify binding partners that interact specifically with phosphorylated Sept7 in order to understand its role in dendritic spine development. Among the proteins we identified, we focus on the 14-3-3 family which are highly conserved phosphoprotein binding proteins that are important for brain development. We identify a specific interaction of pSept7 with 14-3-3γ (gamma) encoded by *YWHAG*. 14-3-3 gamma is highly expressed during brain development, and mainly in neurons. Our findings show that 14-3-3γ is specifically enriched in the dendritic spines, and that its expression in neurons leads to increased dendritic spine numbers.

14-3-3 proteins carry out a diverse array of functions in cellular processes, including in apoptosis (Masters and Fu, 2001), cell cycle progression (Peng et al., 1997), cytoskeletal rearrangements (Gohla and Bokoch, 2002), and neuronal growth (Cornell and Toyo-oka, 2017) are mediated through the multitude of their phosphorylation dependent protein interactors. Accumulating evidence supports a profound role of 14-3-3 proteins in brain development. 14-3-3 proteins are important for neurogenesis, neuronal differentiation, and neuronal migration during cortical development (Cornell and Toyo-oka, 2017). Functional knockout of 14-3-3 proteins in mice brains with the dimeric 14-3-3 inhibitor Difopein results in reduced dendritic complexity and spine density accompanied by schizophrenia-related behaviors (Foote et al., 2015). 14-3-3 proteins are further required for hippocampal long term potentiation (LTP) and associative learning (Qiao et al., 2014), however, contribution of different 14-3-3 proteins in LTP was not ascertained. 14-3-3γ plays an important role in neuronal migration as either knockdown or overexpression of 14-3-3γ results in neuronal migration defects *in vivo* in mice (Wachi et al., 2015). Notably, 14-3-3γ expression is significantly reduced in human brain from Down Syndrome patients (Peyrl et al., 2002), indicating that perturbation of 14-3-3γ levels could contribute to disorders in human brain development. Further, deletion of the *YWHAG* gene encoding 14-3-3γ is also associated with epilepsy and autistic traits in patients with atypical Williams Beuren syndrome due to deletions in 7q11.23 locus (Fusco et al., 2014).

Septins are important regulator of several aspects of neuronal development including dendrite growth, axon development and dendritic spine maturation. Our data show that phosphorylated Sept7 associates with 14-3-3 proteins. While our mass spectrometry data indicates that isoforms gamma, zeta, and epsilon all bind specifically with phosphorylated Sept7 C-terminal tail, we only were able to biochemically validate the interaction with 14-3-3 gamma. It remains unknown whether different isoforms can heterodimerize to associate with phosphorylated Sept7. Using peptide inhibitor Difopein, we found that blocking interaction of Sept7 with 14-3-3 led to a decrease in the level of phosphorylated Sept7 showing the importance of the interaction. Since Difopein quenches the blocking site for phosphoproteins in all 14-3-3 proteins, our study did not directly test isoform specific effect of blocking Sept7 interaction with 14-3-3. Contribution of phosphorylated Sept7 and hence its protection by 14-3-3 proteins in these diverse neuronal contexts will be an important area of study. Dysfunction in both 14-3-3 and septins have been associated with various diseases including neurodegenerative diseases (Foote and Zhou, 2012) (Marttinen et al., 2015) (Ide and Lewis, 2010). Further understanding the interplay of septins with 14-3-3 proteins in the nervous system may provide important insights into the pathophysiology underlying these diseases.

## Acknowledgements

We are grateful for research funding provided by National Institute of Mental Health, R00MH108648 to SY. Mass spectrometry was supported by instrumentation and funding to SEO by R01GM129090. We thank Dr. Yi Zhou for generously sharing the EYFP-Difopein and its negative control expression constructs.

## Author Contributions

SB and BW contributed equally to the work including neuronal studies, biochemical assays, data analyses and manuscript preparation. RF performed neuronal experiments, biochemical assays and molecular biology experiments. KD and CS performed peptide pulldown and mass spectrometry experiments. SEO supervised the mass spectrometry and MS data analyses. SY designed and supervised all experiments, obtained funding and wrote the manuscript.

## REFERENCES

Ageta-Ishihara, N., Miyata, T., Ohshima, C., Watanabe, M., Sato, Y., Hamamura, Y., Higashiyama, T., Mazitschek, R., Bito, H., and Kinoshita, M. (2013). Septins promote dendrite and axon development by negatively regulating microtubule stability via HDAC6-mediated deacetylation. Nat Commun 4, 2532.

Anda, F.C. de, Rosario, A.L., Durak, O., Tran, T., Gräff, J., Meletis, K., Rei, D., Soda, T., Madabhushi, R., Ginty, D.D., et al. (2012). Autism spectrum disorder susceptibility gene TAOK2 affects basal dendrite formation in the neocortex. Nat. Neurosci. 15, 1022–1031.

Berg, D., Holzmann, C., and Riess, O. (2003). 14-3-3 proteins in the nervous system. Nat Rev Neurosci 4, 752–762.

Cornell, B., and Toyo-oka, K. (2017). 14-3-3 Proteins in Brain Development: Neurogenesis, Neuronal Migration and Neuromorphogenesis. Front Mol Neurosci 10, 318.

Ewers, H., Tada, T., Petersen, J.D., Racz, B., Sheng, M., and Choquet, D. (2014). A Septin-Dependent Diffusion Barrier at Dendritic Spine Necks. PLoS ONE 9, e113916.

Finnigan, G.C., Booth, E.A., Duvalyan, A., Liao, E.N., and Thorner, J. (2015). The Carboxy-Terminal Tails of Septins Cdc11 and Shs1 Recruit Myosin-II Binding Factor Bni5 to the Bud Neck in Saccharomyces cerevisiae. Genetics 200, 843–862.

Finnigan, G.C., Duvalyan, A., Liao, E.N., Sargsyan, A., and Thorner, J. (2016). Detection of protein-protein interactions at the septin collar in Saccharomyces cerevisiae using a tripartite split-GFP system. Mol. Biol. Cell 27, 2708–2725.

Foote, M., and Zhou, Y. (2012). 14-3-3 proteins in neurological disorders. Int J Biochem Mol Biology 3, 152–164.

Foote, M., Qiao, H., Graham, K., Wu, Y., and Zhou, Y. (2015). Inhibition of 14-3-3 Proteins Leads to Schizophrenia-Related Behavioral Phenotypes and Synaptic Defects in Mice. Biol Psychiat 78, 386–395.

Fusco, C., Micale, L., Augello, B., Pellico, M.T., Menghini, D., Alfieri, P., Digilio, M.C., Mandriani, B., Carella, M., Palumbo, O., et al. (2014). Smaller and larger deletions of the Williams Beuren syndrome region implicate genes involved in mild facial phenotype, epilepsy and autistic traits. Eur J Hum Genet 22, 64–70.

Gohla, A., and Bokoch, G.M. (2002). 14-3-3 Regulates Actin Dynamics by Stabilizing Phosphorylated Cofilin. Curr Biol 12, 1704–1710.

Hu, J., Bai, X., Bowen, J.R., Dolat, L., Korobova, F., Yu, W., Baas, P.W., Svitkina, T., Gallo, G., and Spiliotis, E.T. (2012). Septin-driven coordination of actin and microtubule remodeling regulates the collateral branching of axons. Curr. Biol. 22, 1109–1115.

Ide, M., and Lewis, D.A. (2010). Altered cortical CDC42 signaling pathways in schizophrenia: implications for dendritic spine deficits. Biol. Psychiatry 68, 25–32.

Kinoshita, M. (2003a). The septins. Genome Biol 4, 236.

Kinoshita, M. (2003b). Assembly of Mammalian Septins. J Biochem 134, 491–496.

Marques, I. de A., Valadares, N.F., Garcia, W., Damalio, J.C.P., Macedo, J.N.A., Araújo, A.P.U. de, Botello, C.A., Andreu, J.M., and Garratt, R.C. (2012). Septin C-terminal domain interactions: implications for filament stability and assembly. Cell Biochem. Biophys. 62, 317–328.

Marttinen, M., Kurkinen, K.M., Soininen, H., Haapasalo, A., and Hiltunen, M. (2015). Synaptic dysfunction and septin protein family members in neurodegenerative diseases. Mol Neurodegener 10, 16.

Masters, S.C., and Fu, H. (2001). 14-3-3 Proteins Mediate an Essential Anti-apoptotic Signal*. J Biol Chem 276, 45193–45200.

Mendonça, D.C., Guimarães, S.L., Pereira, H.D., Pinto, A.A., Farias, M.A. de, Godoy, A.S. de, Araujo, A.P.U., Heel, M. van, Portugal, R.V., and Garratt, R.C. (2021). An atomic model for the human septin hexamer by cryo-EM. J Mol Biol 433, 167096.

Mostowy, S., and Cossart, P. (2012). Septins: the fourth component of the cytoskeleton. Nat. Rev. Mol. Cell Biol. 13, 183–194.

Nourbakhsh, K., Ferreccio, A.A., Bernard, M.J., and Yadav, S. (2021). TAOK2 is an ER-localized Kinase that Catalyzes the Dynamic Tethering of ER to Microtubules. Biorxiv 2021.04.22.440958.

Peng, C.Y., Graves, P.R., Thoma, R.S., Wu, Z., Shaw, A.S., and Piwnica-Worms, H. (1997). Mitotic and G2 checkpoint control: regulation of 14-3-3 protein binding by phosphorylation of Cdc25C on serine-216. Sci New York N Y 277, 1501–1505.

Peyrl, A., Weitzdoerfer, R., Gulesserian, T., Fountoulakis, M., and Lubec, G. (2002). Aberrant expression of signaling - related proteins 14-3-3 gamma and RACK1 in fetal Down Syndrome brain (trisomy 21). Electrophoresis 23, 152–157.

Qiao, H., Foote, M., Graham, K., Wu, Y., and Zhou, Y. (2014). 14-3-3 Proteins Are Required for Hippocampal Long-Term Potentiation and Associative Learning and Memory. J Neurosci 34, 4801–4808.

Richter, M., Murtaza, N., Scharrenberg, R., White, S.H., Johanns, O., Walker, S., Yuen, R.K.C., Schwanke, B., Bedürftig, B., Henis, M., et al. (2018). Altered TAOK2 activity causes autism-related neurodevelopmental and cognitive abnormalities through RhoA signaling. Mol. Psychiatry 20, 1237.

Sirajuddin, M., Farkasovsky, M., Hauer, F., Kühlmann, D., Macara, I.G., Weyand, M., Stark, H., and Wittinghofer, A. (2007). Structural insight into filament formation by mammalian septins. Nature 449, 311–315.

Tada, T., Simonetta, A., Batterton, M., Kinoshita, M., Edbauer, D., and Sheng, M. (2007). Role of Septin cytoskeleton in spine morphogenesis and dendrite development in neurons. Curr. Biol. 17, 1752–1758.

Wachi, T., Cornell, B., Marshall, C., Zhukarev, V., Baas, P.W., and Toyo-oka, K. (2015). Ablation of the 14-3-3gamma Protein Results in Neuronal Migration Delay and Morphological Defects in the Developing Cerebral Cortex: 14-3-3γ Regulates Neuronal Migration. Dev Neurobiol 76, 600–614.

Walikonis, R.S., Jensen, O.N., Mann, M., Provance, D.W., Mercer, J.A., and Kennedy, M.B. (2000). Identification of proteins in the postsynaptic density fraction by mass spectrometry. J Neurosci Official J Soc Neurosci 20, 4069–4080.

Wang, B., Yang, H., Liu, Y.-C., Jelinek, T., Zhang, L., Ruoslahti, E., and Fu, H. (1999). Isolation of High-Affinity Peptide Antagonists of 14-3-3 Proteins by Phage Display †. Biochemistry-Us 38, 12499–12504.

Xie, Y., Vessey, J.P., Konecna, A., Dahm, R., Macchi, P., and Kiebler, M.A. (2007). The GTP-binding protein Septin 7 is critical for dendrite branching and dendritic-spine morphology. Curr. Biol. 17, 1746–1751.

Yadav, S., Oses-Prieto, J.A., Peters, C.J., Zhou, J., Pleasure, S.J., Burlingame, A.L., Jan, L.Y., and Jan, Y.-N. (2017). TAOK2 Kinase Mediates PSD95 Stability and Dendritic Spine Maturation through Septin7 Phosphorylation. Neuron 93, 379–393.

